# The LH receptor regulates hippocampal spatial memory and restores dendritic spine density in ovariectomized APP/PS1 AD mice

**DOI:** 10.1101/2023.12.22.573087

**Authors:** Megan Mey, Sabina Bhatta, Sneha Suresh, Luis Montero Labrador, Helen Piontkivska, Gemma Casadesus

**Author notes:** Corresponding Author: Gemma Casasdesus, Address: Phone Number.

## Abstract

Activation of the luteinizing hormone receptor (LHCGR) rescues spatial memory function and spine density losses associated with gonadectomy and high circulating gonadotropin levels in females. However, whether this extends to the AD brain or the mechanisms that underlie these benefits remain unknown. To address this question, we delivered the LHCGR agonist human chorionic gonadotropin (hCG) intracerebroventricularly (ICV), under reproductively intact and ovariectomized conditions to mimic the post-menopausal state in the APP/PS1mouse brain. Cognitive function was tested using the Morris water maze task, and hippocampal dendritic spine density, Aβ pathology, and signaling changes associated with these endpoints were determined to address mechanisms. Here we show that central LHCGR activation restored function in ovariectomized APP/PS1 female mice to wild-type levels without altering Aβ pathology. LHCGR activation increased hippocampal dendritic spine density regardless of reproductive status, and this was mediated by BDNF-dependent and independent signaling. We also show that ovariectomy in the APP/PS1 brain elicits an increase in peripherally derived pro-inflammatory genes which are inhibited by LHCGR activation. This may mediate reproductive status specific effects of LHCGR agonism on cognitive function and BDNF expression. Together, this work highlights the relevance of the LHCGR on cognition and its therapeutic potential in the “menopausal” AD brain.

## 1. Introduction

Menopausal elevations in circulating gonadotropins are clinically linked to cognitive dysfunction and increased AD risk in women (Butchart et al., 2013; Hogervorst et al., 2004; Hyde et al., 2010; Short et al., 2001; Verdile et al., 2014; Xiong et al., 2022) and in rodent models of menopause (Barron et al., 2010; Berry et al., 2008; Burnham et al., 2017) or high gonadotropins (Casadesus et al., 2007; Xiong et al., 2022). This is further substantiated by rodent studies demonstrating cognitive improvements when elevated circulating gonadotropin levels in ovariectomized mice are pharmacologically lowered (Blair et al., 2016; Bryan et al., 2010; Casadesus et al., 2007; Palm et al., 2014; Telegdy et al., 2009; Ziegler & Thornton, 2010). Notably, this extends to females in AD mouse models (Palm et al., 2014) but not males (Otto et al., 2010), a finding mirrored in human AD patients (Bowen et al., 2015). These functional improvements are paralleled by cellular signaling cascades that positively regulate cognition and neuronal plasticity processes (Blair et al., 2016, 2019; Emanuele et al., 1981; Palm et al., 2014). However, the exact mechanisms underlying gonadotropin effects on the CNS remain poorly understood.

The expression of the luteinizing hormone receptor (LHCGR) (Apaja et al., 2004; Lei et al., 1993; Liu et al., 2007; Yang et al., 2007; Ryu et al., 2022) and LH (Blair et al., 2016, 2019; Emanuele et al., 1981; Palm et al., 2014) have been reported in the central nervous system (CNS), including the hippocampus. Within the hippocampus, CNS LHCGR activation elicits electrophysiological changes (Gallo et al., 1972) that have been directly linked to the regulation of hippocampal-dependent behavior (Yang et al., 2007; Mak et al., 2007) and neurogenesis (Mak et al., 2007). Furthermore, its activation is known to drive signaling cascades associated with the formation and maintenance of hippocampal-dependent memories (Ascoli et al., 2002; Gudermann et al., 1992; Herrlich et al., 1996; Lei & Rao, 2001; Zhang et al., 1999), suggesting that LHCGR activation can positively modulate hippocampal function. Importantly, we (Blair et al., 2019) and others (Emanuele et al., 1981; Glass & McClusky, 1987) have reported that the expression of LH within the brain is inverse to systemic levels. This inverse relationship was also observed in ovariectomized 3xTg-AD female mice under pharmacological inhibition of systemic gonadotropin levels (Palm et al., 2014), which improved cognitive function and positively correlated to CNS LH levels. Therefore, while the central and peripheral administration of high doses of LHCGR ligands such as hCG have been reported to negatively impact CNS function (Burnham et al., 2017) and cause sedation (Lukacs et al., 1995), low doses of centrally administered hCG improve cognition and rescue dendritic spine plasticity in ovariectomized female mice (Blair et al., 2019). These functional and plasticity benefits mimic those observed by downregulation of systemic gonadotropins (Blair et al., 2016) and estradiol therapy in ovariectomized rodents (Gould et al., 1990; Woolley & McEwen, 1992), which also lowers systemic gonadotropins through negative feedback loop (Klein, 2003). The mechanism underlying the reversal between central and systemic LH levels or how circulating LH inhibition drives impoved function is unknown. However, it suggests a more complex signaling mechanism at hand that simply high levels of LH crossing the blood brain barrier (BBB) and persistently driving the LHCGR, as previously suggested (Bowen et al., 2015; Kumar, 2007; Lin et al., 2010; Telegdy et al., 2009; Ziegler & Thornton, 2010).

Therefore, to more directly evaluate the fundamental role of the LHCGR in hippocampal function and dendritic spine plasticity and address its therapeutic potential in AD, we addressed whether centrally delivered hCG could improve cognitive and spine density losses in intact and ovariectomized APP/PS1 AD mice. We also evaluated spatial memory function under hippocampal LHCGR knockdown and determined potential signaling mechanisms, namely BDNF, that link LHCGR signaling to these functional and cellular endpoints and carried out full transcriptome analysis to identify potential mechanistic targets.

## 2. Methods

### 2.1 Animals

To determine the effects of LHCGR signaling on cognitive function, dendritic spines, and related signaling in menopause model AD mice, female (B6.Cg Tg(APPswe, PSEN1dE9) 85Dbo/Mmjax) (APP/PS1) (n=38) were randomly assigned to 4 groups and compared to wild-type sham-operated littermate mice (*Table 1A)*. A second cohort of 21 healthy 6-month-old cycling female (C57/BJ6) mice, had their estrous cycle tracked by vaginal cytology for 4 weeks to determine relationships between estrous cycle hormone changes and signaling proteins (*Table 1B*). All mice were within 2 weeks of age of each other and group-housed in accordance with Kent State University and University of Florida Institutional Animal Care and Use Committee. Food and water were provided ad libitum during a 12-hour light: dark cycle. At the end of the study, mice were deeply anesthetized and sacrificed for tissue collection.

**Table 1:** Treatment groups with initial sample size number. (A) Initial sample size for animal cohort used to determine the effect of LHCGR signaling on cognitive function in menopause model AD mice. (B) Initial sample size for animal cohort used to evaluate serum LH in cycling females with signaling proteins in the hippocampus.

|  |  |  |
| --- | --- | --- |
| **A.** | **Treatment Group** | **Initial Sample Size** |
|  | ADTg+OVX+aCSF | 10 |
|  | ADTg+OVX+hCG | 10 |
|  | ADTg+aCSF | 10 |
|  | ADTg+hCG | 8 |
|  | WT | 13 |
| **B.** | **Estrous Cycle Phase** | **Initial Sample Size** |
|  | Proestrus | 7 |
|  | Estrus | 4 |
|  | Metestrus | 5 |
|  | Diestrus | 5 |

### 2.2 Bilateral Ovariectomy

Animals underwent either bilateral ovariectomy or sham surgery at approximately 6 months old, prior to overt Aβ pathology in the APP/PS1 mouse model. Ovariectomies were conducted as described previously in (Blair et al., 2016). Briefly, animals were deeply anesthetized with 3% isoflurane gas, and ovaries were bilaterally removed through a single dorsal incision or exposed but not excised and reinserted into the abdominal cavity (sham). Wound clips were used to seal the incision and mice were placed in clean cages to recover. Mice received pre and post-surgical pain medication and were monitored daily until wound clips were removed after 10 days.

### 2.3 Cannula implantation

Two months following ovariectomy surgery, at around 8 months old, mice underwent stereotaxic surgery to implant a brain infusion cannula (Alzet; Brain Infusion Kit 3) in the right lateral ventricle using the following coordinates from bregma; anterior posterior −.05mm, medial/lateral −0.11, and dorsal ventral −.25mm. This delay between ovariectomy and initiation of treatment delivery ensured that brain LH levels lowered and stabilized post-ovariectomy as previously determined in (Blair et al., 2019). Drug delivery was performed by connecting the cannula to a subcutaneously placed osmotic pump (Alzet; #1004) filled with hCG (30 mIU/day [Sigma Aldrich, MO]) or artificial cerebrospinal fluid (aCSF) as vehicle. Osmotic pumps were replaced once, after 4 weeks. Cannula placement was verified by injecting fast green through the tubing at sacrifice, only animals with functional attached tubing shown by presence of fast green in the ventricle were included in the study.

### 2.4 Vaginal Cytology & hormone measurements

Estrous cycle was tracked daily at 10:00 am by vaginal lavage with sterile saline. Samples were analyzed daily using a brightfield Olympus microscope at 10X magnification. Estrous cycle stage was determined by the ratio of leukocytes, cornified epithelial cells, and nucleated epithelial cells as described in (Cora et al., 2015).

All blood serum hormone measurements were carried out by the University of Virginia (UVA) Ligand Core. Samples were prepped and serum was collected following their protocols. Briefly, blood was left to clot at room temperature for 90 minutes, spun down at 2000g for 15 minutes to collect serum prior to shipping to UVA for LH measurements and human chorionic gonadotropin (hCG) using the following assays respectively; LH Ultrasensitive – mouse and rat (UVA in house protocol) and HCG – Human Chorionic Gonadotropin (IMMULITE 2000). Results located in supplemental materials (*Supplemental Figure 1*).

### 2.5 Morris Water Maze Testing

Spatial learning and memory was measured using Morris water maze task. A platform was placed in the target quadrant of the pool (120 cm wide) and submerged under opaque white water at 0.5cm deep. The water was temperature controlled at 24°C. Mice were given an initial habituation trial, 24hr prior to onset of training sessions, to familiarize them with the water. Training consisted of 4 trials per day for 5 days. Each trial was started by placing the mice in one of the four quadrants of the pool facing the wall. Sessions were timed out at 60 seconds or when the animal reached the platform. All trials were tracked using a Noldus Ethovision system (Noldus Information Technology Inc., VA). On the 5^th^ day, trial 4 was used as a probe trial where the platform was removed and the mice were allowed to swim for 60 seconds. At the completion of probe trial testing, all mice underwent a visible platform test to verify visual acuity. This involved placing the mouse in the pool, this time with the platform visible and being moved to all 4 quadrants. Only animals that swam to the platform in under 20 seconds in four consecutive trials or whose average was less than 30 seconds after 8 trials were included in the data analysis.

### 2.6 Novel Environment/Open Field Behavior

In the cohort used to determine the effect of LHCGR signaling on cognitive function in menopause model AD mice, general activity and habituation were measured using the Open Field test as described in (Blair et al., 2019). Briefly, mice were placed in a brightly lit open field for 15 minutes. A four-unit open field was used allowing all mice from one cage to be run at one time. Movement and velocity were tracked and time spent in the center, middle, and outer zone was monitored using Noldus Ethovision system (Noldus Information Technology Inc., VA). The trial was analyzed both in totality and in 5-minute blocks to monitor habituation and zone preference.

### 2.7 Light/Dark Box

Anxiolytic behavior was measured using the Light/Dark box procedure. The apparatus (H: 40cm, W: 20cm, L: 40) used for this task had a fully lit chamber (H: 40cm, W: 10cm, L: 40) and a dark chamber (H: 40cm, W: 10cm, L: 40) connected by a 3cm x 5cm opening. The trial had a 5-minute duration beginning with the placement of the animal in the fully lit side of the apparatus. Animals were able to move freely between the light and dark chambers. The number of crosses between chambers and the time spent in each chamber was recorded by the experimenter. Frequency of entries into and the percent time spent in the lit space were analyzed to indicate bright-space anxiety. Results located in supplemental materials (*Supplemental Figure 3*).

### 2.8 Thioflavin S imaging

The left hemisphere was preserved in 4% PFA at 4°C overnight and placed in 30% sucrose until sunk. Brains were flash-frozen in OCT and cut at 40um thickness using a cryostat. Starting at the beginning of the hippocampus, [−.94mm from Bregma], every 6^th^ slice was stained using 1% ThioflavinS. Sections were taken through a series of 5-minute incubation steps at room temperature, as described in (Corrigan et al., 2023). Sections were then mounted on slides, allowed to dry, then taken through EtOH and Xylene dehydration and clearing steps and coverslipped using permount mounting medium (Electron Microscopy Science, PA). A Leica DMi8 automated microscope with Thunder 3D assay imaging software (Leica Microsystems, IL) was used to capture, stitch, and compress Z stack images of entire brain sections at 40X magnification ImageJ was used to quantify plaque number and percent plaque area in the cortex and hippocampus of each tissue section. The data from each tissue section was averaged for each animal.

### 2.9 Soluble Aβ40 and Aβ42 measurements

Cortical tissue was used to assess the amount of soluble Aβ_40_ and Aβ_42_. Hippocampal tissue was not available for this assay because it was used for biochemical analyses. Soluble Aβ measurements were adapted from (Head et.al., 2010). Briefly, tissue was first homogenized in 1X PBS + 200mg/mL protease inhibitor cocktail and spun down at 20,800g for 30 minutes at 4°C. Supernatant was collected as the PBS fraction and stored at −80°C until use. A small amount of the PBS soluble fraction was set aside for protein assay to determine total protein. After the PBS fraction supernatant was removed the pellet was resuspended in 2% SDS + 200mg/mL protease inhibitor cocktail, sonicated, and spun down at 20,800g for 30 minutes at 4°C. The supernatant was collected as SDS soluble fraction and saved at −80°C until use. The remaining pellet was resuspended in 70% formic acid + 200mg/mL protease inhibitor cocktail, sonicated, and spun down at 20,800g for 30 minutes at 4°C. The supernatant was collected as formic acid soluble fraction and was neutralized with 1M Tris (pH 11.0) at a 1:20 ratio. Once all fractions were collected, Human Aβ40 and Aβ42 levels were measured using commercial kits (Invitrogen, MA). Assay values were normalized to total protein.

### 2.10 Golgi Staining & dendritic spine analysis

The left hemisphere of mice sacrificed for dendritic spine density analysis was dissected fresh and immediately underwent the Golgi staining process using the FD rapid Golgi stain kit (FD NeuroTechnologies, MD) per manufacturer’s protocol. All brains were then flash frozen in OCT solution and cut at 100um thickness using a cryostat (Leica CM 1950) (Leica Biosystems, Illinois/USA). Pyramidal neurons in dorsal CA1 region of the hippocampus, extending from 2.7mm to 4.6mm posterior to bregma (Craig et al., 2020), were imaged using the Leica DMi8 automated microscope (Leica Microsystems, IL) with a motorized stage, 63X oil objective. Neurons were selected based on the following criteria (1) the dendrite of interest was isolated from other neurons and dendrites, and (2) the entirety of the neuron was fully impregnated (Craig et al., 2020). Approximately 10-12 neurons in total per animal taken from 3 separate sections, were analyzed and averaged together. The free ImageJ plugin, SpineJ, was used to analyze spines from compressed Z-stacks taken at .27um intervals. The apical dendrite was analyzed on each neuron starting about 10um from the soma. Dendritic spine density was calculated and reported as spines per 10um.

### 2.11 Western Blot Protein Measurements

Hippocampal tissue was homogenized fresh after dissection using cell lysis buffer (Cell Signaling Technology, MA). Protein samples were separated using SDS-PAGE 10% precast gels (Bio-Rad, CA), and transferred to a PVDF membrane. 5% percent milk or bovine serum albumin (depending on antibody manufacturer’s suggestion) was used to block the membranes for one hour at room temperature. Membranes were incubated in primary antibodies overnight in 1X TBST at 4°C: BDNF (1:2,000; Santa Cruz Biotechnology, Texas/USA), Rac1 (1:1,000; Cell Signaling Technology, MA), TrkB (1:1,000; Proteintech, IL), TrkB ser478 (1:1,000; Biosensis, CA), LHCGR (1:500; Proteintech, IL), and actin (1;50,000; Millipore, MA). Membranes were washed and incubated in secondary antibody with horseradish peroxidase for 1 hour at room temperature. Signal was detected by incubation with ECL (Millipore, MA) and imaged with the GBox mini-imaging system (Syngene; MD). ImageJ software (NIH) was used to calculate optical density differences.

### 2.12 Statistical Analysis

All animals were age-matched and randomized to treatment groups. Experimenters were blinded to all treatment groups for all experiments. Analysis was performed using GraphPad Prism 9.4.1 software (Dotmatics, MA). All data passed normality and homogeneity of variance testing so parametric statistics were utilized. Repeated measures ANOVA was used to analyze Morris water maze training. All other analyses were performed as one-way ANOVAs with Tukey post-hoc comparisons, or independent samples t-test where two groups were compared.

### 2.13 Transcriptome sequencing and differential gene expression

Hippocampi were collected and sent out for RNA sequencing to determine effects of hCG on the transcriptome. 1 μg total of RNA was sent to the Case Western Reserve Genomics Core for RNA-sequencing. RNA quality was analyzed using QuBit and only samples that yielded a RIN score of ≥ 8.5 were used to make RNA libraries and later used for sequencing. Transcriptomes were sequenced using Illumina NextSeq with 75 bp single-end reads at a sequencing depth of ∼40 million reads on average per sample. Reads were mapped to the mouse mm38 genome assembly using STAR (ref). Differentially expressed genes DEGs were identified using cuffdiff ver. 2.2.1 (ref), using Fragments Per Kilobase of transcript per Million mapped reads (FPKM) as a gene expression measure normalized to take into account gene length and the total number of mapped reads. False-discovery rate (FDR) based on Benjamini-Hochberg correction (ref) cut-off of 0.05 was used. We further narrowed down the list of candidate DEGs by setting padj value to a stricter 0.05 cut-off, together with a fold change (FC) cut-off of 1.5, a FC suggested to be biologically relevant [81,82].

### 2.14. Pathway enrichment analyses and visualization

To better understand the biological significance of identified DEGs, functional enrichement analysis of DEGs, we used advanced pathway and gene analysis using i-phathway software (Advaitia, CA) and the open software platform, gprofiler (version e110_eg57_p18_4b54a898) (Tartu University, Estonia). Bonferroni correction for multiple tests was used for statistical enrichment cut-off threshold (p-adjusted value of 0.05). Morpheus open software (Broad Institute, MA) was used to produce the heatmap comparison across groups.

## 3. Results

### 3.1 Spatial memory performance in Morris Water Maze in hCG treated AD-Tg mice

Spatial learning and memory differences were measured using Morris Water Maze training and probe trial. All animals passed the visual test. One-way repeated measures ANOVAs were used to determine differences in escape latency, path length, and velocity over training days. Analysis for escape latency showed a significant within-subjects difference for ‘day’ (F(4,44) = 17.88, p < 0.0001) indicating learning across days regardless of treatment group, a between-subjects ‘treatment group’ difference (F(4,44) = 3.075, p < 0.05), indicating treatment differences, and the interaction between ‘day’ and ‘treatment group’ showed a strong trend toward significance (F(4,44) = 1.644, p = 0.063). Individual one-way ANOVAs were performed on each day of training to determine differences across groups during individual days. On day 5, there was a significant difference in escape latency between groups (F(4,44) = 8.716, p <0.0001). Tukey post hoc analysis showed a significantly faster escape latency in the hCG-treated OVX APP/PS1 compared to the sham APP/PS1 mice (p <0.05) and OVX APP/PS1 mice (p <0.001). Similarly, the wild-type control group also showed a significantly faster escape latency compared to the sham APP/PS1 mice (p <0.05) and to OVX APP/PS1 mice (p <0.0001). The hCG-treated sham APP/PS1 mice showed no differences compared to any other groups indicating no cognitive improvements (*Figure 2A*). Analysis for path length showed a significant within-subjects difference for ‘day’ (F(4,44) = 9.113, p < 0.0001) indicating learning across days regardless of treatment group, no between-subjects ‘treatment group’ difference (F(4,44) = 0.6490, p > 0.05), but a significant interaction between ‘day’ and ‘treatment group’ (F(4,44) = 1.720, p < 0.05), indicating a treatment effect dependent on day. Individual one-way ANOVAs were performed on each day of training to determine path length differences across groups during individual days. On day 5, there was a significant difference for path length between groups (F(4,44) = 4.286, p <0.01). Tukey post hoc analysis showed a significantly shorter path length in the hCG-treated OVX APP/PS1 compared to the OVX APP/PS1 mice (p <0.05). Similarly, the wild-type control group also showed a significantly shorter path length than OVX APP/PS1 mice (p <0.01). Analysis for velocity showed a significant within-subjects difference for ‘day’ (F(4,44) = 13.80, p < 0.0001) indicating a change in velocity across days regardless of treatment group, no between-subjects ‘treatment group’ difference (F(4,44) = 2.141, p > 0.05), and no significant interaction between ‘day’ and ‘treatment group’ (F(4,44) = 1.16, p > 0.05), indicating no overall treatment differences in velocity. Altogether, this data illustrates that hCG treatment in OVX APP/PS1 mice normalizes task learning to WT control levels.

**Figure 1:**
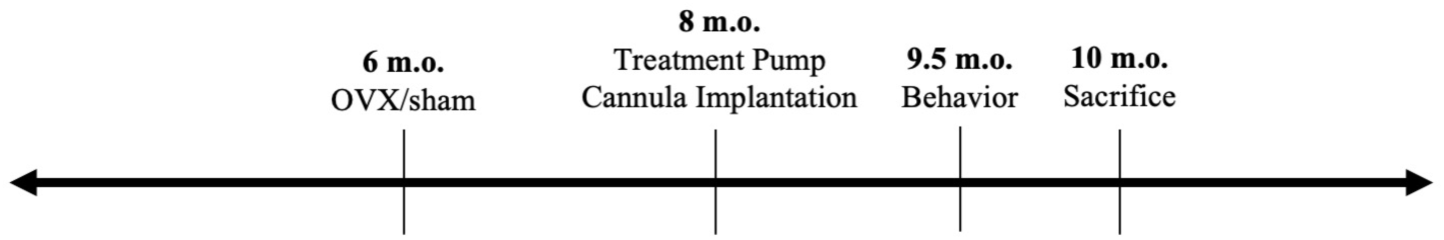
Experimental timeline of surgeries and behavior in AD-Tg cycling and OVX mice treated ICV with aCSF or hCG.

**Figure 2.**
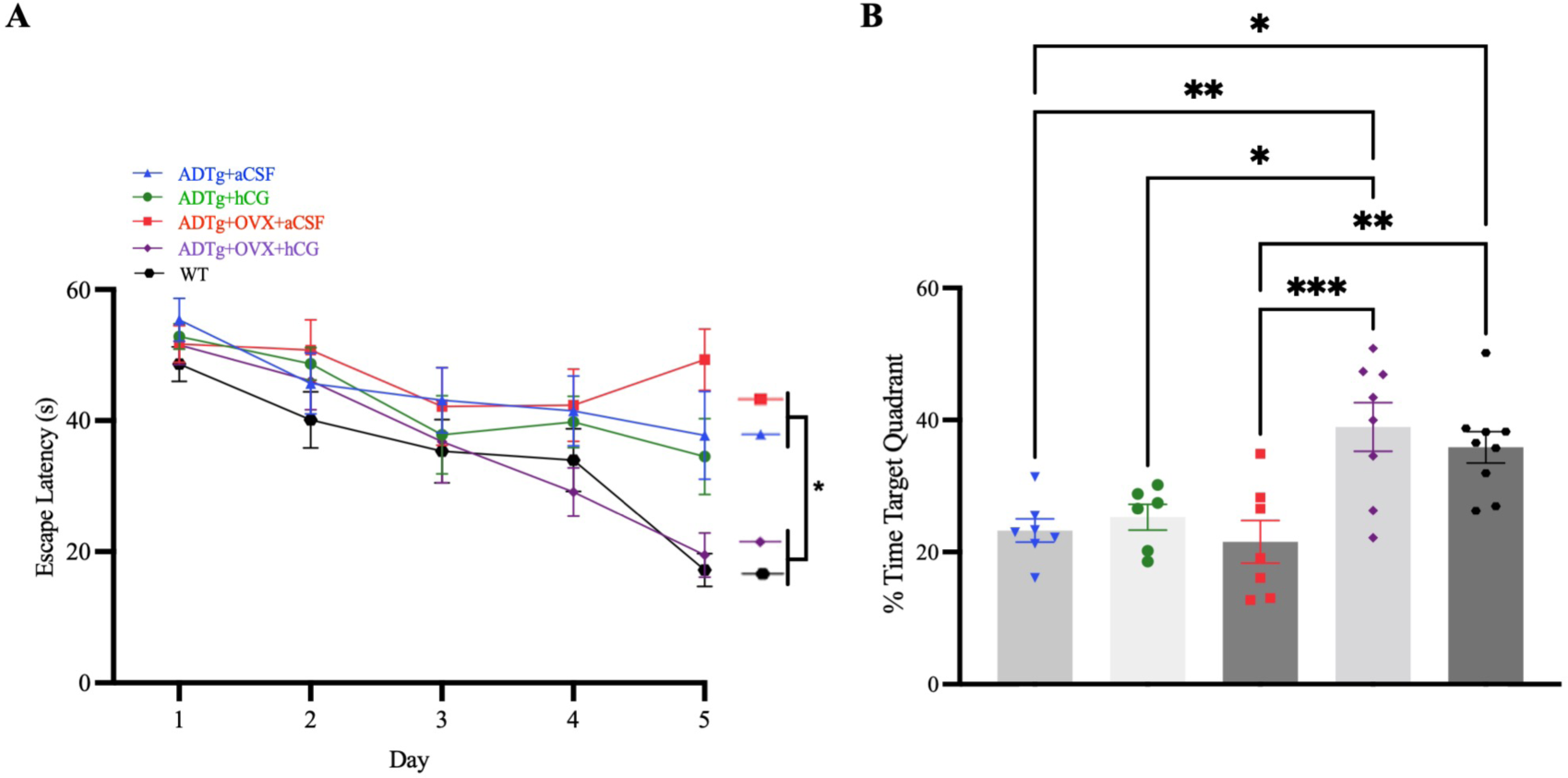
Morris Water Maze performance. (A) Training: Escape latency per training day represented as mean ± SEM per group (ADTg+aCSF, n=10; ADTg+hCG, n=8; ADTg+OVX+aCSF, n=8; ADTg+OVX+hCG, n=10; WT Control, n= 13). (B) Probe trial: Percent time in the target quadrant represented as mean ± SEM per group (ADTg+aCSF, n=7; ADTg+hCG, n=6; ADTg+OVX+aCSF, n=7; ADTg+OVX+hCG, n=8; WT Control, n= 9). Significant differences denoted as *p<0.05, ** p<0.01, *** p<0.001.

Spatial memory retention group differences were determined by the probe trial on trial 4 of day 5 of training and were analyzed using a one-way ANOVA. Due to COVID-19 water shut-offs a portion of the last cohort of animals in this study endured colder-than-usual water during probe trial. Due to the distress this caused, animals had to be removed prior to the end of the trial and thus have been removed probe trial analysis (APP/PS1+aCSF n=3, APP/PS1+hCG n=2, APP/PS1 OVX+aCSF n=3, and APP/PS1 OVX+hCG n=2). However, despite the smaller number of animals that completed the probe trial in each group, our data show a significant difference across groups in percent-time spent in the target quadrant (F(4,32) = 8.238, p < 0.0001). Tukey post hoc analysis revealed that the hCG-treated OVX APP/PS1 mice spent significantly more time in the target quadrant compared to the sham APP/PS1 mice (p = .0031), hCG-treated sham APP/PS1 (p <0.05), and OVX APP/PS1 animals (p <0.01). The WT control group spent significantly more time in the target quadrant compared to sham APP/PS1 (p <0.01) and OVX APP/PS1 (p <0.01) mice. Once again, the hCG-treated sham APP/PS1 mice showed no differences compared to other groups indicating no cognitive improvements. There were no differences between hCG-treated OVX APP/PS1 and WT control mice, again demonstrating that hCG treatment in OVX APP/PS1 mice normalized spatial memory retention to WT mice (*Figure 2B*).

### 3.2 Aβ plaque deposition & soluble Aβ40 and Aβ42 measurements

Thioflavin S staining was carried out to determine the impact of ovariectomy and/or treatment on dense core Aβ plaque deposits in the cortex and hippocampus of all APP/PS1 treatment groups. One-way ANOVA analysis showed no significant differences between groups in total Aβ plaque count in the cortex or hippocampus (*Figure 3A-B*).

**Figure 3.**
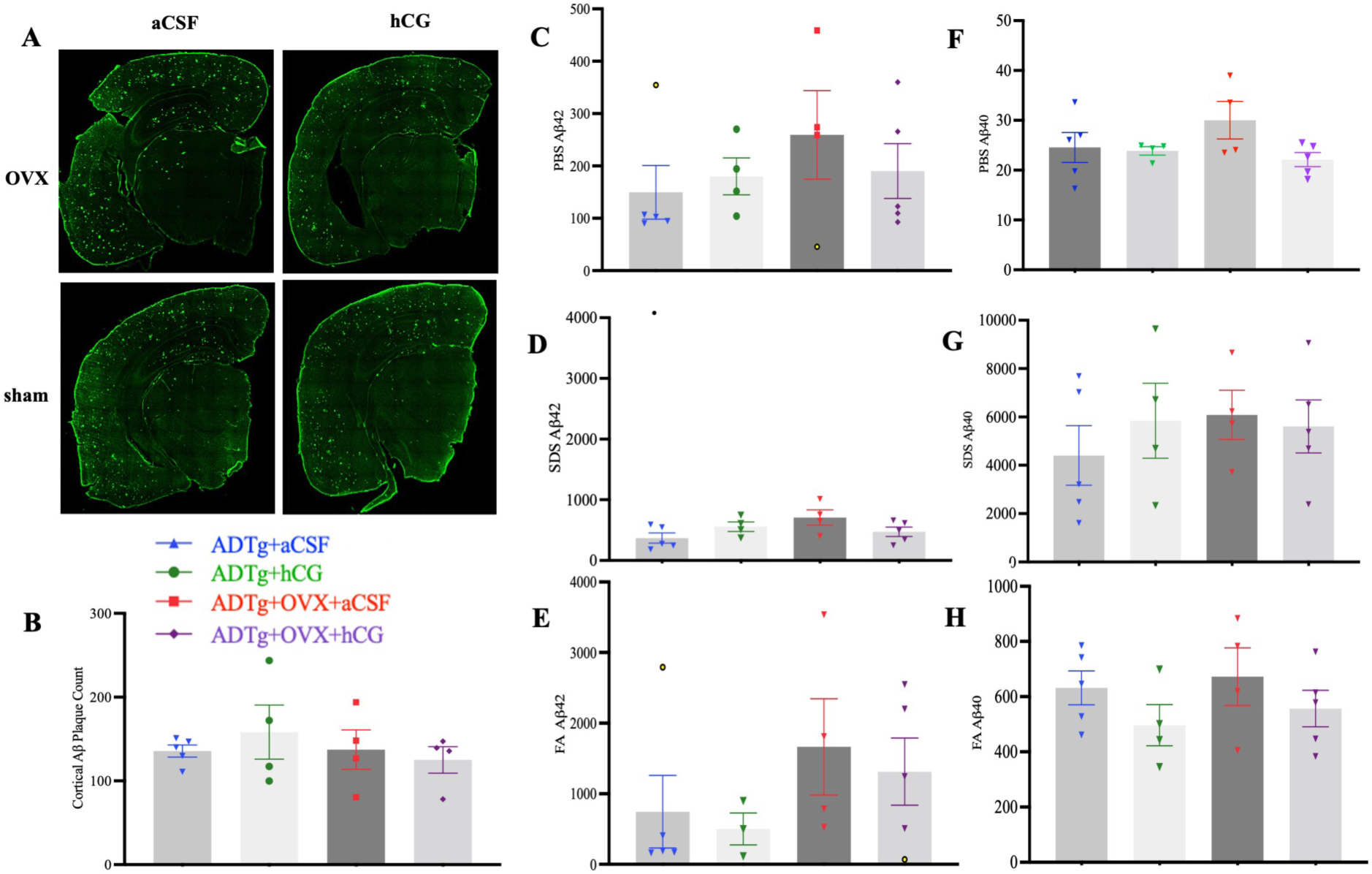
Cortical amyloid-β pathology. (A) Representative images of thioflavin stained sections for each group (B) Cortical total thioflavin-A positive plaque count. (C - E) Cortical Aβ42 soluble fractions. (F - H) Cortical Aβ40 soluble fractions. Data represented as mean ± SEM for each measure per group (ADTg+aCSF, n=5; ADTg+hCG, n=4; ADTg+OVX+aCSF, n=4; ADTg+OVX+ hCG, n=4). Significant differences denoted as *p<0.05, ** p<0.01, *** p<0.001.

Cortical PBS, SDS, and formic acid soluble Aβ_40_ and Aβ_42_ fractions were also analyzed using a one-way ANOVA. There were no significant differences in any of the Aβ soluble forms across treatment groups (*Figure 3C-H*).

### 3.5 Dendritic spine density analysis

To evaluate the effect of hCG treatment on dendritic spine density in our treatment groups we counted spines in the CA1 region of the hippocampus because of its involvement in spatial navigation and memory (Alexander et al., 2020), and the expression of LHCGR in this region (Ryu et al., 2022). One-way ANOVA and Tukey’s post hoc analysis were used to compare average spines per 10um between groups in each region. Spine density within the CA1 region showed significant differences across treatment groups (F(4,16) = 9.190, p <0.001). Aligned with the cognitive improvements, the hCG-treated OVX APP/PS1 mice showed higher spine density compared to the sham APP/PS1 mice (p <0.01) and OVX APP/PS1 mice (p <0.05). Interestingly, although they did not show cognitive improvements, hCG-treated sham APP/PS1 mice also had significantly higher spine densities compared to the sham APP/PS1 mice (p <0.05) and OVX APP/PS1 mice (p <0.05). Lastly, the wild-type control group showed higher spine densities compared to sham APP/PS1 mice (p <0.01) and OVX APP/PS1 mice (p <0.01) and were not significantly different compared to hCG-treated sham or OVX APP/PS1 mice (*Figure 4*). Together, these data show that hCG treatment increased dendritic spine density in the CA1 region of the hippocampus back to control WT levels regardless of reproductive status.

**Figure 4.**
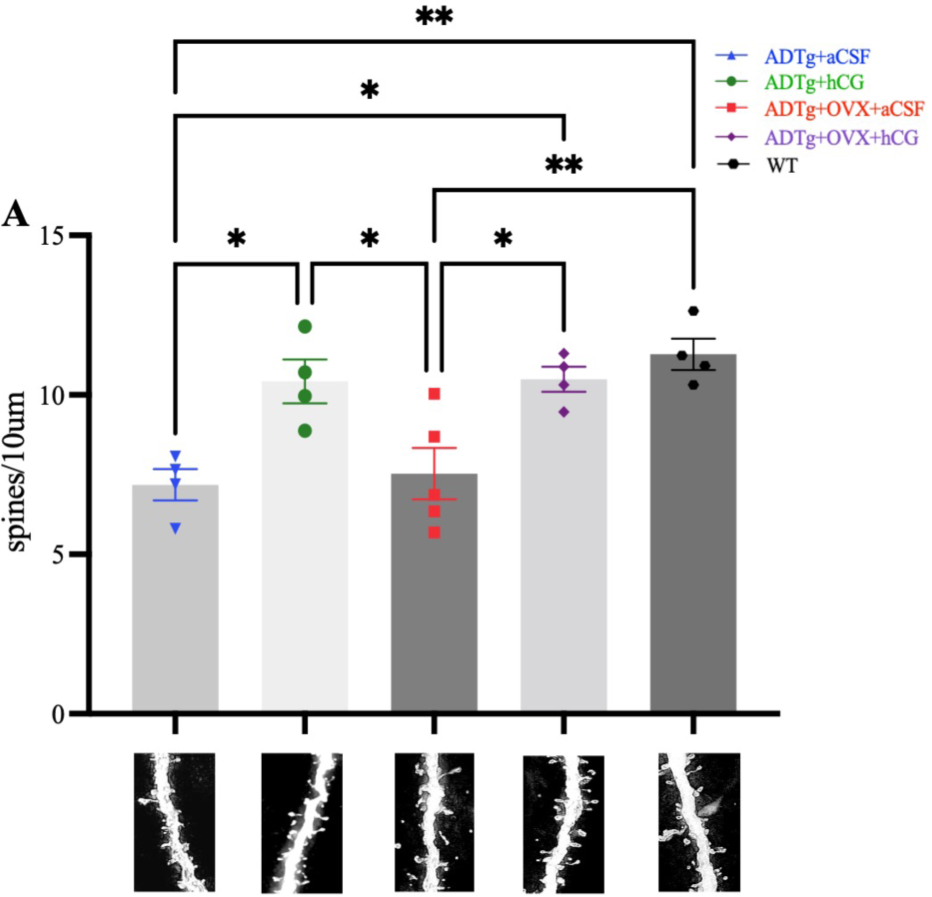
Golgi stain of Dendritic spines in the hippocampal CA1 region. Dendritic spines per 10µm, represented as mean ± SEM per group (ADTg+aCSF, n=4; ADTg+hCG, n=4; ADTg+OVX+aCSF, n=4; ADTg+OVX+hCG, n=4; WT Control n= 4). Significant differences denoted as *p<0.05, ** p<0.01.

### 3.6 Western Blot protein measurements

BDNF is involved in cognition, neuroplasticity, and dendritic spine remodeling (Miranda et al., 2019 for review). Therefore, we measured BDNF in the hippocampus. We also determined signaling protein changes that are downstream of BDNF and involved in dendritic spine development including the BDNF receptor TrkB phosphorylation at serine 478, and the downstream spine elongation factor Rac1. To this end, one-way ANOVA analysis revealed no significant differences in proBDNF. However, there were significant differences across groups in hippocampal mature BDNF (F(4,18) = 4.333, p <0.05). Aligned with the cognitive and dendritic spine density data, tukey post hoc analysis showed that the hCG-treated OVX APP/PS1 mice showed significantly higher BDNF expression compared to the OVX APP/PS1 mice (p <0.05). Wild-type control group also showed significantly higher BDNF expression compared to OVX APP/PS1 mice (p <0.05). The hCG-treated sham APP/PS1 mice did not show any significant differences in mature BDNF, indicating that dendritic spine density was not modulated by BDNF in this group (*Figure 5A*).

**Figure 5.**
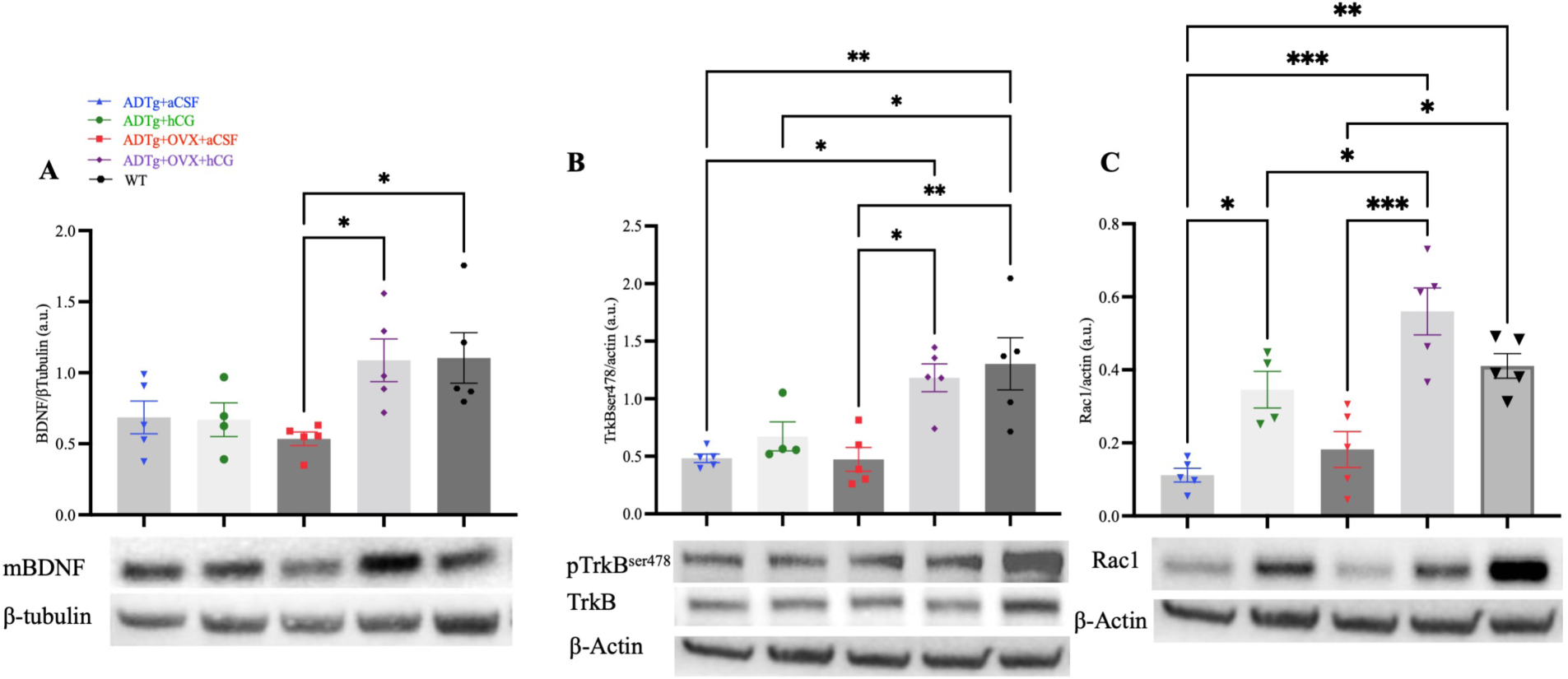
Dendritic spine morphogenesis-related signaling in the hippocampus. (A) Mature BDNF expression (16kDa) and β-Tubulin (55kDa), (B) TrkB and phosphorylated TrkB^ser478^ (90kDa) and β-actin (42kDa) and (C) Rac1 (21kDa) and β-actin (42kDa) represented as mean ± SEM for each protein per group (ADTg+aCSF, n=5; ADTg+hCG, n=4; ADTg+OVX+aCSF, n=5; ADTg+OVX+hCG, n=5; WT Control, n= 5). Significant differences denoted as *p<0.05, ** p<0.01, *** p<0.001.

Similar to BDNF, there were also significant differences in expression of its receptor TrkB phosphorylated at serine 478 (F(4,19) = 8.350, p <0.001), which is directly involved in mediating dendritic spine morphology (Gough, 2012; Lai et al., 2012). Tukey post hoc comparisons showed that the hCG-treated OVX APP/PS1 mice had significantly higher relative TrkBser478 expression levels compared to OVX APP/PS1 mice (p <0.05). The WT control group also had significantly higher relative TrkB ser478 expression compared to OVX APP/PS1 mice (p <0.05). The hCG-treated sham APP/PS1 mice did not show any significant differences compared to any other group (*Figure 5B*). This further supports a potential role of BDNF signaling in the cognitive improvements and increased dendritic spine density in the hCG treated OVX APP/PS1 group.

The dendritic spine elongation factor, Rac1, which lies downstream of TrkB serine 478 phosphorylation also showed significant group differences (F(4,19) = 15.96, p <.0001). Tukey post hoc analysis showed significantly higher expression of Rac1 in the hCG-treated OVX APP/PS1 compared to the sham APP/PS1 mice (p <.001), OVX APP/PS1 mice (p <.001) and hCG-treated sham APP/PS1 (p <0.05). Interestingly, the hCG-treated sham APP/PS1 mice, although not as high as OVX hCG-treated mice, showed significantly higher relative expression of Rac1 compared to the sham APP/PS1 mice (p <0.05), possibly accounting for the dendritic spine increases observed in this group. Lastly, wild-type control mice had higher relative expression of Rac1 compared to the sham APP/PS1 mice (p < 0.001) and OVX APP/PS1 mice (p < 0.01) (*Figure 5C*). Lastly, because of the sensitivity of LHCGR to ligand concentration we measured LHCGR expression in the hippocampus and found no differences in expression between groups (*Supplemental Figure 4B*).Together, these data indicate that hCG regulates signaling events that are directly linked to spine development highlighting the trophic potential and a novel mechanism associated with LHCGR activation in the CNS.

We also measured mature hippocampal BDNF in a separate cohort of healthy cycling WT mice to evaluate the relationship between serum LH across the estrus cycle and hippocampal BDNF. One-way ANOVA showed a significant difference between mice sacrificed during different phases of the estrous cycle, F(3,17) = 4.750, p <0.01. Tukey post hoc analysis showed significantly higher hippocampal BDNF during estrus compared to proestrus (p<0.05), metestrus (p<0.05) and diestrus (p<0.05) (*Figure 6A*). There were no significant differences in blood serum LH between groups (*Figure 6B*). However, there was a significant negative correlation between serum LH and mature hippocampal BDNF expression; p<0.05, R=-.365 (*Figure 6C*). This indicates that the fluctuation of LH across the estrous cycle mirror fluctuations in hippocampal BDNF, as reported for estrogen (Gibbs, 1998), and suggests a potential contribution of LH in menstural-cycle associated cognitive and plasticity differences. Of note, lowering post-ovariectomy rises in peripheral LH, which restores CNS LH levels (Blair et al., 2019; Emanuele et al., 1981; Palm et al., 2014), restores cognition (Bohm-Levine et al., 2020; Bryan et al., 2010; Palm et al., 2014; Ziegler & Thornton, 2010), recovers ovariectomy-associated spine density (Blair et al., 2019) and also increases BDNF trancription (Palm et al., 2014) and protein expression (Bohm-Levine et al., 2020). Together with the above findings, our data supports a connection between LH signaling and BDNF expression.

**Figure 6.**
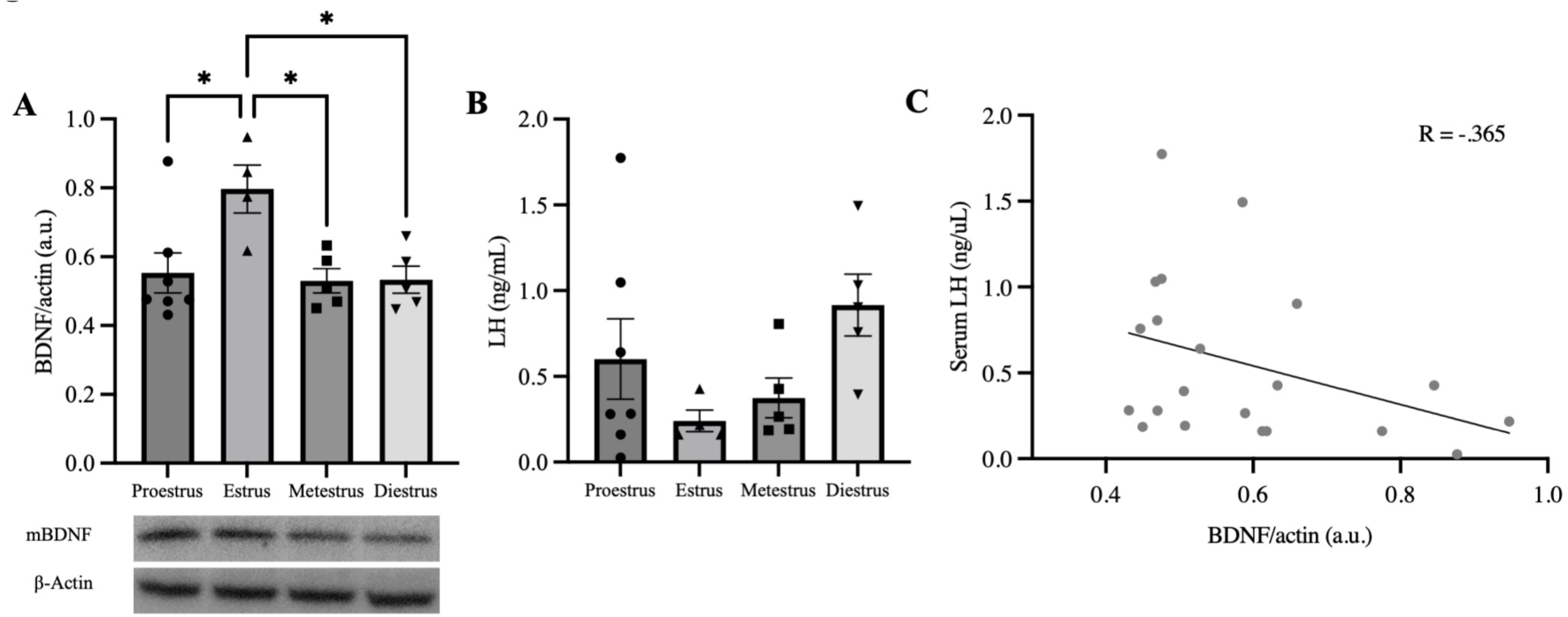
Relationship between serum LH and BDNF levels. (A) mature hippocampal BDNF (16kDa) and β-actin (42kDa) and (B) Serum LH levels (ng/ml) in cycling female WT mice. Data is represented as mean SEM per estrus cycle time (proestrus, n=7; estrus, n=4; metestrus, n=5; and diestrus, n=5). (C) Pearson’s correlation between serum LH and hippocampal mBDNF. Significant differences denoted as *p<0.05.

### 3.7 Differential gene expression due to OVX and hCG treatments

To identify genes and pathways that could be contributing to the observed functional improvements in OVX APP/PS1 but not in intact APP/PS1 mice, we have examined hippocampus transcriptomes using bulk RNA sequencing (Figure 7). When intact APP/PS1 mice were compared with OVX and intact animals receiving hCG, there were 163 and 106 DEGs respectively. Comparisons between OVX and OVX+hCG treatments yielded 147 DEGs. Notably, 20 genes overlapped between the these comparisons which were evaluated further in the context of potential mechanism (Figure 7A). Functional annotation of these 20 DEGs identified the top 5 overrepresented GO terms and pathways, as immune system related (Figure 7B). Heat mapping of these genes revealed that OVX significantly increased such genes compared to intact APP/PS1 mice. hCG treatment in intact APP/PS1 mice lead to changes similar to those observed by OVX, suggesting that LHCGR signaling in intact rodents mimics ovariectomy/menopausal status effects. However, hCG treatment in OVX mice led to an inhibition of these genes compared to WT control levels (Figure 7C), suggesting that reproductive status regulates the ability of hCG to modulate inflammatory signals. A deeper dive into genes that were changed by hCG in the OVX condition revealed that a significant number of genes upregulated by OVX or treatment of hCG in intact APP/PS1 animals that were normalized by hCG in the ovariectomized group were immune related and linked to peripheral myeloid cell lineage and CNS infiltration (Figure 7D). Furthermore, our analysis designed to identify the top 5 upstream genes linked the observed DEGs yielded the three G-protein signaling modulators (GPSM1, 2 &3), Purkinje cell protein 2 (PCP2), and protein tyrosine phosphatase non-receptor type 6 all noted immune function regulators.

**Figure 7.**
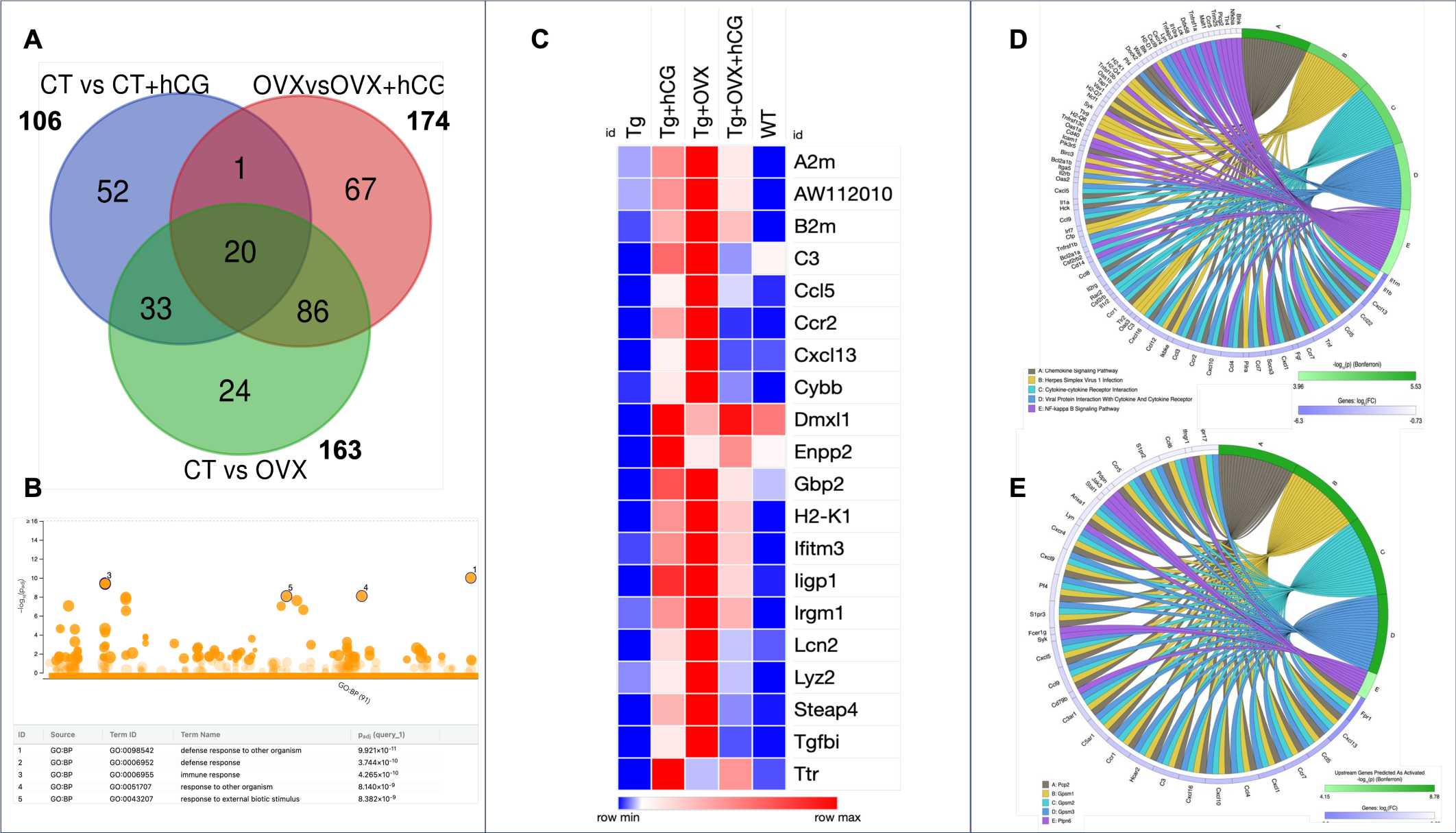
**(A)** differentially expressed genes numbers shared between three pairwise comparisons in APP/PS1 transgenics (ADtg), under intact, ovariectomized and hCG treated conditions (ADTg+aCSF vs ADTg+OVX+aCSF, ADTg+aCSF vs ADTg+hCG, and ADTg+OVX+aCSF vs ADTg+OVX+hCG). (B) Top 5 Go Terms (biological Process) identified using GProfiler functional analysis of 20 overlaping genes across the three DEG comparisons with bonferroni correction and significance cut-off threshold of 0.05 per g:SCS algorithm. **(C)** Heatmap of average expression levels in each APP/PS1 surgery and treatment group and. WT (n=3-4/group)of 20 overlapping genes across three comparisons. **(D)** i-pathway functional pathway analysis of top 50 DEGs identified in the ADtg OVX vs OVX+hCG comparison using a bonferroni correction and significance cut-off threshold of 0.05. **(E)** i-pathway upstream regulator analysis identifying top 5 potential regulators of DEGs identified in the ADtg OVX vs OVX+hCG comparison using a bonferroni correction and significance cut-off threshold of 0.05.

## 4. Discussion

Spatial memory loss is a hallmark of AD (King et al., 2019 for review) and readily detectable in models of menopause (Chesler & Juraska, 2000; Heikkinen et al., 2004; Li et al., 2004; Ziegler & Thornton, 2010) and AD (Knafo et al., 2009; Perez-Cruz et al., 2011; Walker & Herskowitz, 2021). Our results demonstrate that activation of the CNS LHCGR using one of its two ligands, hCG, recovered learning and memory function to wild-type control levels. Notably, benefits of hCG were restricted to our ovariectomized cohort, suggesting that, in cycling animals, gonadal steroids may be working to inhibit the LHCGR signaling benefits, at least in the context of this AD mouse model. It is also important to note that although including reproductively intact hCG-treated AD-Tg mice is an important experimental control, it is less translationally relevant since sporadic AD only occurs in post-menopausal women. Together, this interesting finding highlights the need to account for the complex interactions between gonadal steroids and other HPG axis hormones when addressing the role of reproductive hormones in the aging/AD brain.

Previous work from our laboratory (Blair et al., 2016; Bryan et al., 2010) and that of others (Telegdy et al., 2009; Ziegler & Thornton, 2010) has shown that lowering high post-ovariectomy circulating LH levels improves learning and memory in ovariectomized rodents. This suggests that LHCGR signaling negatively impacts cognition. However, we (Blair et al., 2019; Palm et al., 2014) and others (Emanuele et al., 1981) have shown an inverse relationship between brain and peripheral LH levels and have shown that normalizing LHCGR signaling in conditions of elevated LH improves function and stimulates dendritic spines. This aspect is also supported by correlational findings positively linking CNS LH mRNA expression to improved spatial ability in animals under pharmacological gonadotropin inhibition (Palm et al., 2014).

Notably, others have shown that hCG treatment negatively impacts spatial ability (Barron et al., 2010; Berry et al., 2008; Burnham et al., 2017; Lukacs et al., 1995). These effects, however, were observed under estrogen replacement or in gonadally intact conditions which better mirror our hCG-treated gonadally intact group which did not show benefits of hCG. The lack of effectiveness in this group may highlight pharmacokinetic events associated with hCG dose relative to basal circulating levels of LH (both bind the LHCGR). Our study did not show differences in LHCGR expression levels in any of our hCG treated groups, however the LHCGR is highly sensitive to post-translational modifications, internalization, and transcriptional downregulation, in the presence of excess ligands, or low levels of circulating ligands (Hu et al., 1990, Segaloff et al., 1990, Peegel eta al., 1994, Min et al., Kishi et al., 2001), which could impact LHCGR signaling and drive cognitive deficits. This will need to be addressed in more detail in future studies. Also of note, higher doses of hCG show CNS depressant effects (Lukacs et al., 1995). As shown in our supplementary data, we did not detect any differences in anxiety or locomotor activity in hCG treated animals in this current study or our prior work using low doses (Blair et al., 2019).

Learning and memory performance is intimately linked to the process of dendritic spine plasticity (Frank et al., 2018; Lai et al., 2012; Xiong et al., 2013). Here we report that central delivery of low dose of hCG increases dendritic spines in the CA1 region of the hippocampus. Dendritic spines in the CA1 region are reduced in AD mouse models (Frank et al., 2018; Perez-Cruz et al., 2011; Rodriguez et al., 2013), including APP/PS1 mice (Knafo et al., 2009), and also after ovariectomy (Woolley et al., 1990). Recovery of dendritic spines in this area has been directly linked to improvements in cognitive function (Rodriguez et al., 2013). This suggests, at least partially, that a potential key mediator of the observed spatial memory improvements in our ovariectomized hCG-treated mice is the rescue of dendritic spines in this region.

Interestingly, hCG treatment in reproductively intact mice also led to increases in dendritic spines in the CA1 region of the hippocampus even though these mice did not show cognitive improvements, which illustrates a more complex mechanistic dynamic. To address this paradox, we determined changes in cellular signaling mechanisms associated with spine formation and known to be linked to LHCGR signaling (Kawamura et al., 2005). Here we report that hippocampal mature BDNF levels and the phosphorylation of the BDNF receptor TrkB, known to be necessary for BDNF-driven spine morphogenesis (Gough et al., 2012), were restored to wild-type levels in ovariectomized hCG-treated mice, but not in hCG-treated reproductively intact mice. There is a well established link between LHCGR activation and BDNF regulation in the ovary (Kawamura et al., 2005), thus here we extend this link to the brain, through the ability of LHCGR activation to increase BDNF expression.

Notably, The lack of BDNF differences in hCG-treated cycling animals, again suggests an inhibitory role of gonadal steroids in this system that will be important to mechanistically tease apart. In connection with these findings, we also show that circulating LH levels during the estrus cycle are negatively associated with hippocampal mature BDNF protein expression. This inverse relationship is recapitulated in studies where high circulating gonadotropins were pharmacologically inhibited leading to increases in BDNF levels (Bohm-Levine et al., 2020). It is also important to note that LHCGR (Gómez Ravetti et al., 2010) and LHβ transcription levels (Palm et al., 2014) are reduced in human AD brains which parallel low BDNF levels (Caffino et al., 2020 for review). Together our findings support the involvement of LHCGR in central BDNF regulation and the need to mechanistically evaluate this inverse relationship in the context of local and circulating steroids to fully understand HPG axis hormone influences on learning and memory and AD pathogenesis.

hCG treatment in ovariectomized APP/PS1 mice increased phosphorylation of the TrkB receptor at serine 478. TrkB phosphorylation at this site is essential in mediating spine morphogenesis signaling. Rac1 activation is also vital in this process (Borin et al., 2018; Bosco et al., 2009; Lai & Ip, 2013), and dysregulated in human AD brains (Borin et al., 2018; Kikuchi et al., 2020). In this context, we did not determine the upstream mechanisms that could account for increased TrkB phosphorylation in our animals. CDK5, which is activated by BDNF, is a key kinase that needs future evaluation. Nevertheless, we did address downstream signaling effects of TrkB phosphorylation on Rac1 expression and demonstrate that APP/PS1 mice, under intact and ovariectomized conditions, have significantly reduced levels of Rac1 that are restored by hCG regardless of reproductive status. As expected based on the function of Rac1, these increases mirrored increases in dendritic spines observed in both hCG treated groups. However, given that only ovariectomized-hCG-treated animals showed BDNF/TrkB signaling changes and cognitive improvements, these data suggest a decoupling of BDNF/TrkB, Rac1 regulation, and spine formation that will need to be further explored. Furthermore, our results also suggest that increased spine number in the absence of BDNF/TrkB changes is not sufficient to improve function. This is well described in the literature (Zagrebelsky et al., 2020 for review) and associated with differences in spine subtypes, which will also need to be determined in future work. However, together, our BDNF/TrkB signaling and Rac1 findings explain the decoupling between cognitive benefits and dendritic spine density in our reproductively intact mice.

Of note, hCG treatment improved cognitive function and dendritic spine density independently of Aβ pathology regulation. Our findings contrast previous work showing reduced AD pathology and cognitive improvements in a global embryonic LHCGR knockout-APP^SW+^ cross (Lin et al., 2010). These animals show profound disruptions in gonadal steroid levels, thus a direct link between our findings and this study is difficult to make. However, the neuroprotective effects of hCG in this study, in the presence of high Aβ levels, underscores the importance of BDNF regulation as the main protective mechanism underlying the therapeutic benefits of hCG. This does not exclude other mechanisms of action, especially since hCG is also known for its powerful immune regulating role in the periphery (van Groenendael et al., 2019; Schumacher & Zenclussen, 2019). Based on this, it will become important to address other potential neuroprotective mechanisms of this hormone.

Lastly, in pregnancy hCG is implicated in supporting implantation and placentation by regulating local and maternal innate and adaptive immune responses (Schumacher & Zenclussen, 2017). Our gene and pathway discovery analyses demonstrate than hCG therapeutic effects in ovariectomized APP/PS1 mice may be driven through such immune regulating functions. Interstingly, these effects are only evident in OVX mice, which show a high degree of peripheral immune genes within the hippocampus. LHCGR agonism within the brain of ovariectomized females drives a marked anti-inflammatory response, which our data suggests is mainly related to it’s ability to mitigate peripheral infiltration mechanisms than are accelerated by ovariectomy. The reason why LHCGR activation yields differential responses depending on reproductive status is unknown, and as discussed above, likely to involve a mechanistic relationship with gonadal steroids and or changes in LHCGR surface expression that will need to be uncovered. However, our findings linking LHCGR activation toinhibition of myeloid and hematopoetic immune genes is exciting given the pathogeneic involvement of peripheral pro-inflammatory mechanisms in several CNS conditions and disorders. Notably, G-protein signaling modulators (GPSM1, 2 &3) are receptor-independent G protein signaling regulators with several functions including the myeloid-cell immune regulation (Yan et al., 2022), as is protein tyrosine phosphatase non-receptor type 6 (PTPN6) which is is primarily expressed in hematopoietic cells and a known negative regulator of the immune system (Tsui et al., 2006). Another highlighted upstream regulator, Purkinje cell protein 2 (PCP2), also a G-protein modulator highly expressed in cerebellum, bipolar retinal cells, and coincidently also highly expressed in the testes. PCP2 is known to regulate cell differentiation (Guan et al., 2005) and centrally linked to sensory-motor gating and mood stabilization (Walton et al., 2012). Together these findings demonstrate that at least in this AD mouse model, central LHCGR may be involved in regulating mecahnisms involved in peripheral infiltration and that intermediate mechanisms, namely gonadal steroids, may be interacting with LHCGR signaling.

## 5. Conclusions

CNS gonadotropin receptor signaling has historically (Barron et al., 2010; Berry et al., 2008; Burnham et al., 2017) and more recently (Xiong et al., 2022) been reported to be pathogenic. However, we demonstrate that driving CNS LHCGR activity using low doses of the LH analog hCG restores spatial learning and memory, dendritic spine density, and BDNF signaling, all aspects that are profoundly impacted in AD (Gould et al., 1990; Masliah et al., 2001; Walker & Herskowitz, 2021). We also note that benefits of hCG/LHCGR are independent of changes in Aβ pathology. This suggests that restoration of memory function and dendritic spine density are likely driven through trophic-BDNF or other neuroprotective effects.

Importantly, the benefits of hCG are restricted to the ovariectomized condition, when gonadal steroid levels are low. Our full transcriptome analysis reveals that this may be linked to the ability of LHCGR to regulate the unique pro-inflammatory environment that is facilitated by ovariectomy, namely peripheral infiltration. However, these findings highlight a complex relationship between gonadal steroids and gonadotropin signaling that is currently unexplored and than may help explain the complexity of the effects of steroid therapy on cognition and AD risk in menopausal women. This work also underscores the therapeutic value of targeting this receptor in the menopausal female population.

## Acknowledgements

The University of Virginia Center for Research in Reproduction Ligand Assay and Analysis Core is supported by the Eunice Kennedy Shriver NICHD/NIHGrant R24HD10. This work was supported by NIA R01 funding R01-AG054654-01.

## Supplemental Materials

### Hormone Serum Measurements

To confirm ovariectomy status, we analyzed serum LH levels in sham versus OVX groups (*Supplemental Figure 1*). LH serum levels were significantly different across groups by one-way ANOVA, (F(4,40) = 29.95, p <0.0001). Tukey post hoc analysis demonstrated that OVX APP/PS1 (ADTg+OVX+acSF) mice had significantly elevated serum LH levels compared to sham APP/PS1 mice (ADTg+aCSF) (p <.0001), hCG-treated sham APP/PS1 (ADTg+hCG) (p <.0001), and wild-type littermate (WT) controls (p <0.0001). Similarly, hCG-treated OVX APP/PS1 (ADTg+OVX+hCG) also had significantly elevated serum LH levels compared to all sham groups: APP/PS1(p <.0001), APP/PS1+hCG (p <0.0001), and WT (p <0.0001). There was no effect of hCG treatment on serum LH levels. To identify if the ICV hCG delivery had crossed into the periphery we measured blood serum hCG. There was no detectable hCG in the serum of hCG-treated mice. These data indicate that ovariectomies resulted in the expected rise in LH and that hCG did not cross to the periphery at detectable levels.

**Supplemental Figure 1.**
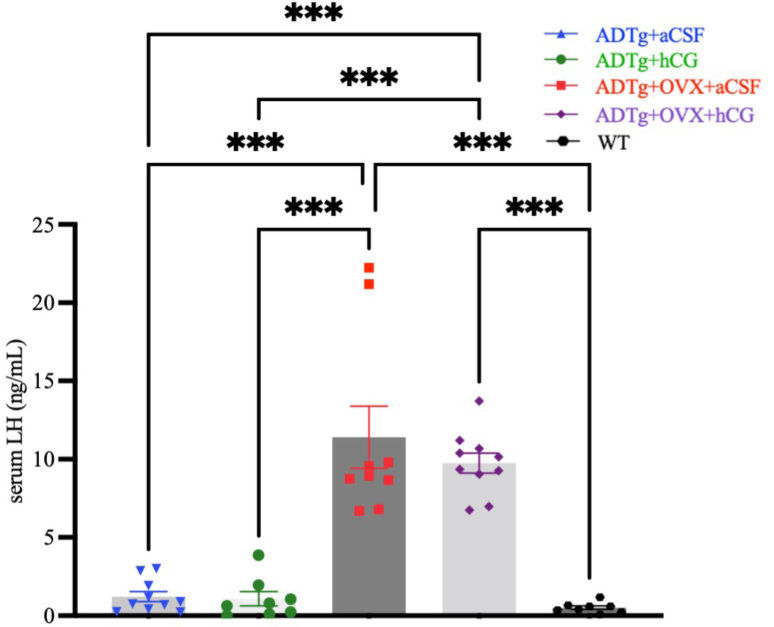
Circulatory LH levels. Serum LH levels (ng/ml) represented as mean ± SEM for each group (ADTg+aCSF, n=10; ADTg+hCG, n=8; ADTg+OVX+aCSF, n=9; ADTg+OVX+hCG, n=10; WT Control n= 8). Significant differences denoted as *p<0.05, ** p<0.01, *** p<0.001.

### Activity, habituation, and anxiolytic behaviors in Open Field and Light/Dark Box

In the cohort used to determine the effect of LHCGR signaling on cognitive function in menopause model AD mice, activity level, habituation, and zone preference were evaluated using the Open Field task. Statistical outliers were removed from the analysis of distance and velocity (APP/PS1 OVX+aCSF n=1). A one-way ANOVA used to analyze total distance, average velocity, and preference for the center or outer zones during the first 5 minutes of the trial. Both total distance and average velocity showed a significant difference between treatment groups, (F(4,45) = 3.82, p < 0.01) and (F(4,45) = 3.83, p < 0.01) respectively. Tukey post hoc analysis showed that sham APP/PS1 mice had increased distance (p < 0.05) and increased velocity (p < .05) compared to WT control animals. This indicates hyperactivity in the APP/PS1 mouse strain, which is a commonly reported phenotype (Huang et al., 2016). There were no significant differences in center or outer zone preference between groups (*Supplemental Figure 2*). A one-way ANOVA was used to compare the ability to habituate across 5 minute time bins between groups. There was no interaction or time bin effect, however there was a a significant effect of treatment group, (F(4,133) = 12.89, p < 0.001. Tukey post analysis showed that only the WT control animals were able to habituate.

Frequency of crosses into the lit chamber of the light/dark box was used analyzed as an indicator of anxiety. Animals that did not cross into the lit chamber were excluded from analysis (APP/PS1+aCSF n=2, APP/PS1 OVX+aCSF n=1, APP/PS1 OVX+hCG n=1, and WT n=1). A one-way ANOVA showed no significant differences between groups (*Supplemental Figure 3*). Altogether this data demonstrates that hCG treatment did not effect activity or anxiety.

**Supplemental Figure 2.**
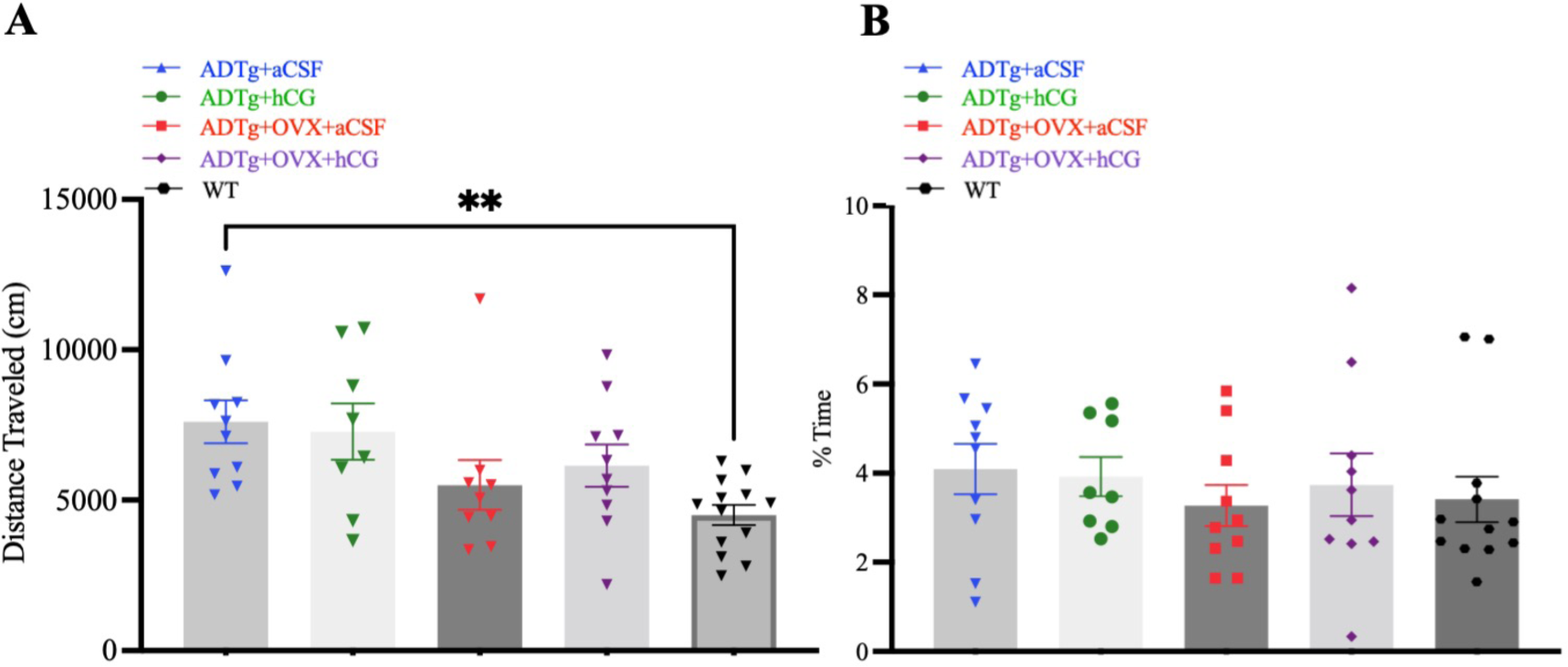
Novel Open Field Activity. Data is represented as mean ± SEM for each group (A) Total distance traveled, (B) percent time spent in the center zone during the first 5 minutes. Data is represented as mean ± SEM for each group (ADTg+aCSF, n=10; ADTg+hCG, n=8; ADTg+OVX+aCSF, n=10; ADTg+OVX+hCG, n=10; WT Control, n= 12). Significant differences denoted as *p<0.05, ** p<0.01.

**Supplemental Figure 3.**
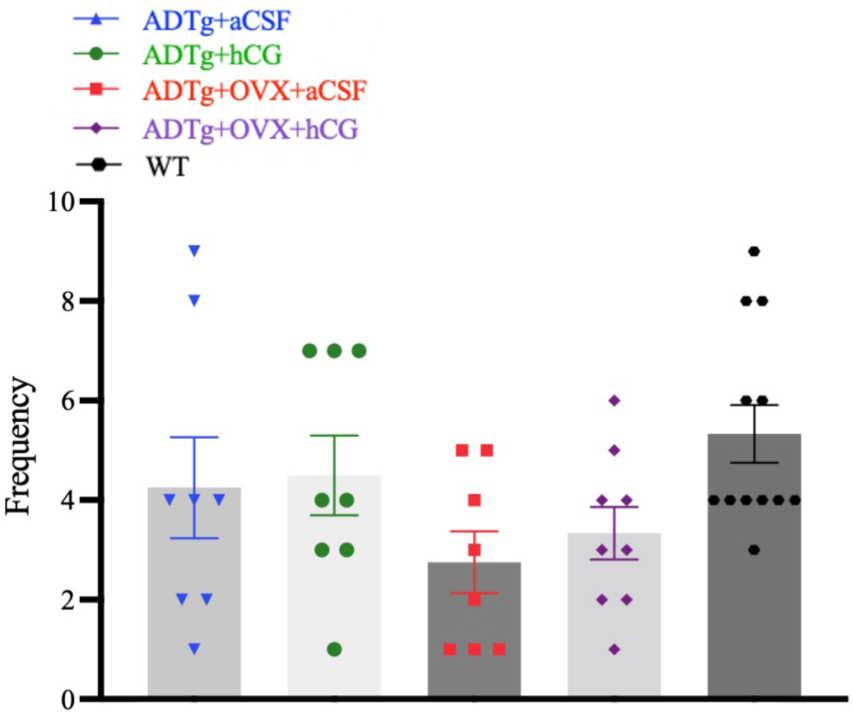
Light/Dark Box Performance. Average cross frequency into the lit chamber ± SEM (ADTg+aCSF, n=8; ADTg+hCG, n=8; ADTg+OVX+aCSF, n=8; ADTg+OVX+hCG, n=9; WT Control, n= 12)

**Supplemental Figure 4.**
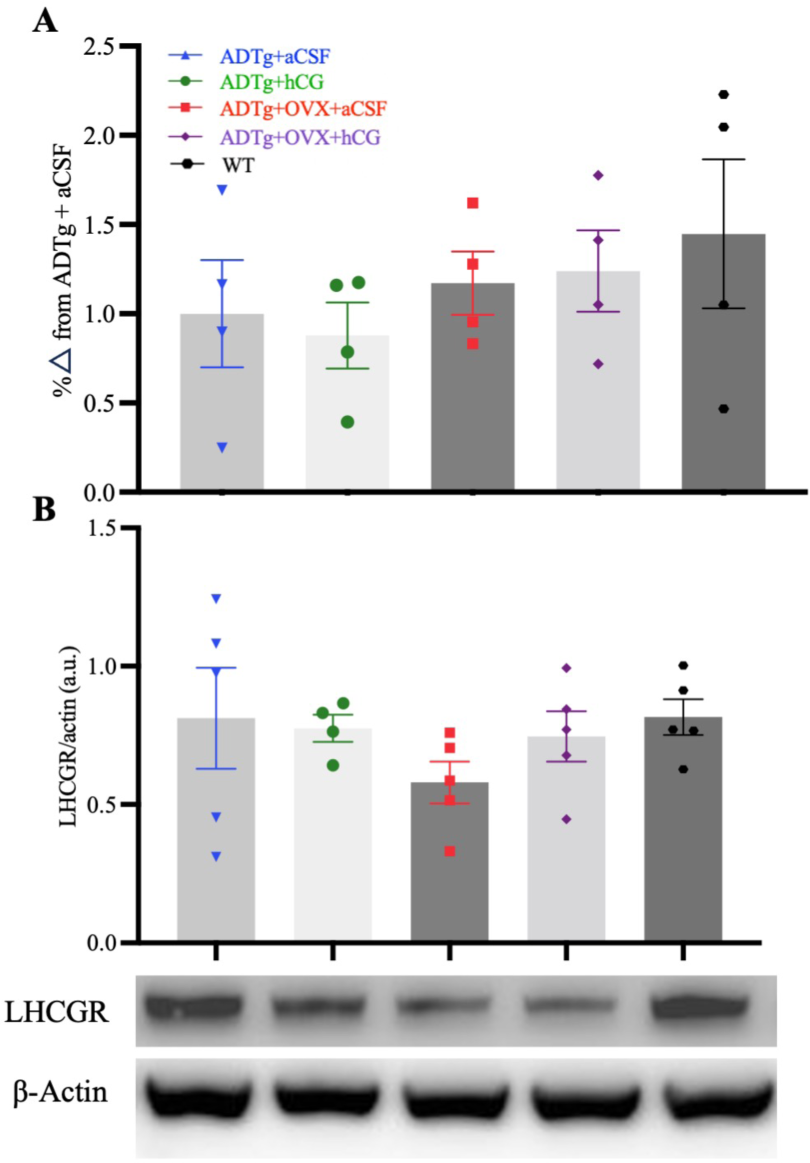
Hippocampal LHCGR transcription and protein expression. Graphs represent mean ± SEM per group **(A)** LHCGR RNA transcripts percent change from control. **(B)** LHCGR protein expression (80kDa) and β-Actin (42kDa), (ADTg+aCSF, n=4; ADTg+hCG, n=4; ADTg+OVX+aCSF, n=4; ADTg+OVX+hCG, n=4; WT Control, n=4).

## References

Apaja, P. M., Harju, K. T., Aatsinki, J. T., Petäjä-Repo, U. E., & Rajaniemi, H. J. (2004). Identification and structural characterization of the neuronal luteinizing hormone receptor associated with sensory systems. The Journal of Biological Chemistry, 279(3), 1899–1906. 10.1074/jbc.M311395200

Ascoli, M., Fanelli, F., & Segaloff, D. L. (2002). The lutropin/choriogonadotropin receptor, a 2002 perspective. Endocrine Reviews, 23(2), 141–174. 10.1210/edrv.23.2.0462

Barron, A. M., Verdile, G., Taddei, K., Bates, K. A., & Martins, R. N. (2010). Effect of Chronic hCG Administration on Alzheimer’s-Related Cognition and Aβ Accumulation in PS1KI Mice. Endocrinology, 151(11), 5380–5388. 10.1210/en.2009-1168

Berry, A., Tomidokoro, Y., Ghiso, J., & Thornton, J. (2008). Human chorionic gonadotropin (a luteinizing hormone homologue) decreases spatial memory and increases brain amyloid-beta levels in female rats. Hormones and Behavior, 54(1), 143–152. 10.1016/j.yhbeh.2008.02.006

Blair, J. A., Bhatta, S., & Casadesus, G. (2019). CNS Luteinizing Hormone Receptor Activation Rescues Ovariectomy-Related Loss of Spatial Memory and Neuronal Plasticity. Neurobiology of Aging, 78, 111–120. 10.1016/j.neurobiolaging.2019.02.002

Blair, J. A., Palm, R., Chang, J., McGee, H., Zhu, X., Wang, X., & Casadesus, G. (2016). Luteinizing hormone downregulation but not estrogen replacement improves ovariectomy-associated cognition and spine density loss independently of treatment onset timing. Hormones and Behavior, 78, 60–66. 10.1016/j.yhbeh.2015.10.013

Bohm-Levine, N., Goldberg, A. R., Mariani, M., Frankfurt, M., & Thornton, J. (2020). Reducing luteinizing hormone levels after ovariectomy improves spatial memory: Possible role of brain-derived neurotrophic factor. Hormones and Behavior, 118, 104590. 10.1016/j.yhbeh.2019.104590

Borin, M., Saraceno, C., Catania, M., Lorenzetto, E., Pontelli, V., Paterlini, A., Fostinelli, S., Avesani, A., Di Fede, G., Zanusso, G., Benussi, L., Binetti, G., Zorzan, S., Ghidoni, R., Buffelli, M., & Bolognin, S. (2018). Rac1 activation links tau hyperphosphorylation and Aβ dysmetabolism in Alzheimer’s disease. Acta Neuropathologica Communications, 6(1), 61. 10.1186/s40478-018-0567-4

Bosco, E. E., Mulloy, J. C., & Zheng, Y. (2009). Rac1 GTPase: A “Rac” of All Trades. Cellular and Molecular Life Sciences : CMLS, 66(3), 370–374. 10.1007/s00018-008-8552-x

Bowen, R. L., Perry, G., Xiong, C., Smith, M. A., & Atwood, C. S. (2015). A clinical study of lupron depot in the treatment of women with Alzheimer’s disease: Preservation of cognitive function in patients taking an acetylcholinesterase inhibitor and treated with high dose lupron over 48 weeks. Journal of Alzheimer’s Disease: JAD, 44(2), 549–560. 10.3233/JAD-141626

Bryan, K. J., Mudd, J. C., Richardson, S. L., Chang, J., Lee, H.-G., Zhu, X., Smith, M. A., & Casadesus, G. (2010). Down-regulation of serum gonadotropins is as effective as estrogen replacement at improving menopause-associated cognitive deficits. Journal of Neurochemistry, 112(4), 870–881. 10.1111/j.1471-4159.2009.06502.x

Burnham, V., Sundby, C., Laman-Maharg, A., & Thornton, J. (2017). Luteinizing hormone acts at the hippocampus to dampen spatial memory. Hormones and Behavior, 89, 55–63. 10.1016/j.yhbeh.2016.11.007

Butchart, J., Birch, B., Bassily, R., Wolfe, L., & Holmes, C. (2013). Male sex hormones and systemic inflammation in Alzheimer disease. Alzheimer Disease and Associated Disorders, 27(2), 153–156. 10.1097/WAD.0b013e318258cd63

Caffino, L., Mottarlini, F., & Fumagalli, F. (2020). Born to Protect: Leveraging BDNF Against Cognitive Deficit in Alzheimer’s Disease. CNS Drugs, 34(3), 281–297. 10.1007/s40263-020-00705-9

Casadesus, G., Milliken, E. L., Webber, K. M., Bowen, R. L., Lei, Z., Rao, C. V., Perry, G., Keri, R. A., & Smith, M. A. (2007). Increases in luteinizing hormone are associated with declines in cognitive performance. Molecular and Cellular Endocrinology, 269(1–2), 107–111. 10.1016/j.mce.2006.06.013

Chesler, E. J., & Juraska, J. M. (2000). Acute Administration of Estrogen and Progesterone Impairs the Acquisition of the Spatial Morris Water Maze in Ovariectomized Rats. Hormones and Behavior, 38(4), 234–242. 10.1006/hbeh.2000.1626

Cora, M. C., Kooistra, L., & Travlos, G. (2015). Vaginal Cytology of the Laboratory Rat and Mouse: Review and Criteria for the Staging of the Estrous Cycle Using Stained Vaginal Smears. Toxicologic Pathology, 43(6), 776–793. 10.1177/0192623315570339

Corrigan, R. R., Labrador, L., Grizzanti, J., Mey, M., Piontkivska, H., & Casadesús, G. (2023). Neuroprotective Mechanisms of Amylin Receptor Activation, Not Antagonism, in the APP/PS1 Mouse Model of Alzheimer’s Disease. Journal of Alzheimer’s Disease: JAD, 91(4), 1495–1514. 10.3233/JAD-221057

Craig, E., Dillingham, C. M., Milczarek, M. M., Phillips, H. M., Davies, M., Perry, J. C., & Vann, S. D. (2020). Lack of change in CA1 dendritic spine density or clustering in rats following training on a radial-arm maze task. Wellcome Open Research, 5, 68. 10.12688/wellcomeopenres.15745.2

Emanuele, N., Oslapas, R., Connick, E., Kirsteins, L., & Lawrence, A. M. (1981). Hypothalamic LH may play a role in control of pituitary LH release. Neuroendocrinology, 33(1), 12–17. 10.1159/000123194

Frank, A. C., Huang, S., Zhou, M., Gdalyahu, A., Kastellakis, G., Silva, T. K., Lu, E., Wen, X., Poirazi, P., Trachtenberg, J. T., & Silva, A. J. (2018). Hotspots of dendritic spine turnover facilitate clustered spine addition and learning and memory. Nature Communications, 9(1), Article 1. 10.1038/s41467-017-02751-2

Gallo, R. V., Johnson, J. H., Kalra, S. P., Whitmoyer, D. I., & Sawyer, C. H. (1972). Effects of luteinizing hormone on multiple-unit activity in the rat hippocampus. Neuroendocrinology, 9(3), 149–157. 10.1159/000122046

Gibbs, R. B. (1998). Levels of trkA and BDNF mRNA, but not NGF mRNA, fluctuate across the estrous cycle and increase in response to acute hormone replacement. Brain Research, 787(2), 259–268. 10.1016/S0006-8993(97)01511-4

Glass, J. D., & McClusky, M. E. (1987). Immunoreactive luteinizing hormone-containing neurons in the brain of the white-footed mouse, Peromyscus leucopus. Experientia, 43(2), 188–190. 10.1007/BF01942846

Gómez Ravetti, M., Rosso, O. A., Berretta, R., & Moscato, P. (2010). Uncovering Molecular Biomarkers That Correlate Cognitive Decline with the Changes of Hippocampus’ Gene Expression Profiles in Alzheimer’s Disease. PLoS ONE, 5(4), e10153. 10.1371/journal.pone.0010153

Gough, N. R. (2012). Connecting TrkB to Dendritic Remodeling. Science Signaling, 5(249), ec287–ec287. 10.1126/scisignal.2003752

Gould, E., Woolley, C. S., Frankfurt, M., & McEwen, B. S. (1990). Gonadal steroids regulate dendritic spine density in hippocampal pyramidal cells in adulthood. The Journal of Neuroscience: The Official Journal of the Society for Neuroscience, 10(4), 1286–1291. 10.1523/JNEUROSCI.10-04-01286.1990.

Guan J, Luo Y, Denker BM. Purkinje cell protein-2 (Pcp2) stimulates differentiation in PC12 cells by Gbetagamma-mediated activation of Ras and p38 MAPK (2005). Biochem J. 392(Pt 2):389–97. doi: 10.1042/BJ20042102.

Gudermann, T., Birnbaumer, M., & Birnbaumer, L. (1992). Evidence for dual coupling of the murine luteinizing hormone receptor to adenylyl cyclase and phosphoinositide breakdown and Ca2+ mobilization. Studies with the cloned murine luteinizing hormone receptor expressed in L cells. The Journal of Biological Chemistry, 267(7), 4479–4488.

Heikkinen, T., Puoliväli, J., & Tanila, H. (2004). Effects of long-term ovariectomy and estrogen treatment on maze learning in aged mice. Experimental Gerontology, 39(9), 1277–1283. 10.1016/j.exger.2004.05.005

Herrlich, A., Kühn, B., Grosse, R., Schmid, A., Schultz, G., & Gudermann, T. (1996). Involvement of Gs and Gi proteins in dual coupling of the luteinizing hormone receptor to adenylyl cyclase and phospholipase C. The Journal of Biological Chemistry, 271(28), 16764–16772. 10.1074/jbc.271.28.16764

Hogervorst, E., Bandelow, S., Combrinck, M., & Smith, A. D. (2004). Low free testosterone is an independent risk factor for Alzheimer’s disease. Experimental Gerontology, 39(11– 12), 1633–1639. 10.1016/j.exger.2004.06.019

Huang, H., Nie, S., Cao, M., Marshall, C., Gao, J., Xiao, N., Hu, G., & Xiao, M. (2016). Characterization of AD-like phenotype in aged APPSwe/PS1dE9 mice. AGE, 38(4), 303–322. 10.1007/s11357-016-9929-7

Hyde, Z., Flicker, L., Almeida, O. P., McCaul, K. A., Jamrozik, K., Hankey, G. J., Chubb, S. A. P., & Yeap, B. B. (2010). Higher luteinizing hormone is associated with poor memory recall: The health in men study. Journal of Alzheimer’s Disease: JAD, 19(3), 943–951. 10.3233/JAD-2010-1342

Kawamura, K., Kawamura, N., Mulders, S. M., Sollewijn Gelpke, M. D., & Hsueh, A. J. W. (2005). Ovarian brain-derived neurotrophic factor (BDNF) promotes the development of oocytes into preimplantation embryos. Proceedings of the National Academy of Sciences of the United States of America, 102(26), 9206–9211. 10.1073/pnas.0502442102

Kikuchi, M., Sekiya, M., Hara, N., Miyashita, A., Kuwano, R., Ikeuchi, T., Iijima, K. M., & Nakaya, A. (2020). Disruption of a RAC1-centred network is associated with Alzheimer’s disease pathology and causes age-dependent neurodegeneration. Human Molecular Genetics, 29(5), 817–833. 10.1093/hmg/ddz320

Klein, C. E. (2003). The Hypothalamic-Pituitary-Gonadal Axis. Holland-Frei Cancer Medicine. 6th Edition. https://www.ncbi.nlm.nih.gov/books/NBK13386/

Knafo, S., Alonso-Nanclares, L., Gonzalez-Soriano, J., Merino-Serrais, P., Fernaud-Espinosa, I., Ferrer, I., & DeFelipe, J. (2009). Widespread changes in dendritic spines in a model of Alzheimer’s disease. Cerebral Cortex (New York, N.Y.: 1991), 19(3), 586–592. 10.1093/cercor/bhn111

Kumar, T. R. (2007). Functional analysis of LHbeta knockout mice. Molecular and Cellular Endocrinology, 269(1–2), 81–84. 10.1016/j.mce.2006.10.020

Lai, K.-O., & Ip, N. Y. (2013). Structural plasticity of dendritic spines: The underlying mechanisms and its dysregulation in brain disorders. Biochimica et Biophysica Acta (BBA) - Molecular Basis of Disease, 1832(12), 2257–2263. 10.1016/j.bbadis.2013.08.012

Lai, K.-O., Wong, A. S. L., Cheung, M.-C., Xu, P., Liang, Z., Lok, K.-C., Xie, H., Palko, M. E., Yung, W.-H., Tessarollo, L., Cheung, Z. H., & Ip, N. Y. (2012). TrkB phosphorylation by Cdk5 is required for activity-dependent structural plasticity and spatial memory. Nature Neuroscience, 15(11), 1506–1515. 10.1038/nn.3237

Lei, Z. M., & Rao, C. V. (2001). Neural Actions of Luteinizing Hormone and Human Chorionic Gonadotropin. Seminars in Reproductive Medicine, 19(1), 103–110. 10.1055/s-2001-13917

Lei, Z. M., Rao, C. V., Kornyei, J. L., Licht, P., & Hiatt, E. S. (1993). Novel expression of human chorionic gonadotropin/luteinizing hormone receptor gene in brain. Endocrinology, 132(5), 2262–2270. 10.1210/endo.132.5.8477671

Li, C., Brake, W. G., Romeo, R. D., Dunlop, J. C., Gordon, M., Buzescu, R., Magarinos, A. M., Allen, P. B., Greengard, P., Luine, V., & McEwen, B. S. (2004). Estrogen alters hippocampal dendritic spine shape and enhances synaptic protein immunoreactivity and spatial memory in female mice. Proceedings of the National Academy of Sciences, 101(7), 2185–2190. 10.1073/pnas.0307313101

Lin, J., Li, X., Yuan, F., Lin, L., Cook, C. L., Rao, C. V., & Lei, Z. (2010). Genetic Ablation of Luteinizing Hormone Receptor Improves the Amyloid Pathology in a Mouse Model of Alzheimer Disease. Journal of Neuropathology & Experimental Neurology, 69(3), 253–261. 10.1097/NEN.0b013e3181d072cf

Liu, T., Wimalasena, J., Bowen, R. L., & Atwood, C. S. (2007). Luteinizing hormone receptor mediates neuronal pregnenolone production via up-regulation of steroidogenic acute regulatory protein expression. Journal of Neurochemistry, 100(5), 1329–1339. 10.1111/j.1471-4159.2006.04307.x

Lukacs, H., Hiatt, E. S., Lei, Z. M., & Rao, C. V. (1995). Peripheral and intracerebroventricular administration of human chorionic gonadotropin alters several hippocampus-associated behaviors in cycling female rats. Hormones and Behavior, 29(1), 42–58. 10.1006/hbeh.1995.1004

Mak, G. K., Enwere, E. K., Gregg, C., Pakarainen, T., Poutanen, M., Huhtaniemi, I., & Weiss, S. (2007). Male pheromone-stimulated neurogenesis in the adult female brain: Possible role in mating behavior. Nature Neuroscience, 10(8), 1003–1011. 10.1038/nn1928

Masliah, E., Mallory, M., Alford, M., DeTeresa, R., Hansen, L. A., McKeel, D. W., & Morris, J. C. (2001). Altered expression of synaptic proteins occurs early during progression of Alzheimer’s disease. Neurology, 56(1), 127–129. 10.1212/wnl.56.1.127

Miranda, M., Morici, J. F., Zanoni, M. B., & Bekinschtein, P. (2019). Brain-Derived Neurotrophic Factor: A Key Molecule for Memory in the Healthy and the Pathological Brain. Frontiers in Cellular Neuroscience, 13. https://www.frontiersin.org/articles/10.3389/fncel.2019.00363

Ondo, J. G., Mical, R. S., & Porter, J. C. (1972). Passage of radioactive substances from CSF to hypophysial portal blood. Endocrinology, 91(5), 1239–1246. 10.1210/endo-91-5-1239

Palm, R., Chang, J., Blair, J., Garcia-Mesa, Y., Lee, H., Castellani, R. J., Smith, M. A., Zhu, X., & Casadesus, G. (2014). Down-regulation of serum gonadotropins but not estrogen replacement improves cognition in aged-ovariectomized 3xTg AD female mice. Journal of Neurochemistry, 130(1), 115–125. 10.1111/jnc.12706

Perez-Cruz, C., Nolte, M. W., Gaalen, M. M. van, Rustay, N. R., Termont, A., Tanghe, A., Kirchhoff, F., & Ebert, U. (2011). Reduced Spine Density in Specific Regions of CA1 Pyramidal Neurons in Two Transgenic Mouse Models of Alzheimer’s Disease. Journal of Neuroscience, 31(10), 3926–3934. 10.1523/JNEUROSCI.6142-10.2011

Rodriguez, G. A., Burns, M. P., Weeber, E. J., & Rebeck, G. W. (2013). Young APOE4 targeted replacement mice exhibit poor spatial learning and memory, with reduced dendritic spine density in the medial entorhinal cortex. Learning & Memory, 20(5), 256–266. 10.1101/lm.030031.112

Ryu, V., Gumerova, A., Korkmaz, F., Kang, S. S., Katsel, P., Miyashita, S., Kannangara, H., Cullen, L., Chan, P., Kuo, T., Padilla, A., Sultana, F., Wizman, S. A., Kramskiy, N., Zaidi, S., Kim, S.-M., New, M. I., Rosen, C. J., Goosens, K. A., … Zaidi, M. (2022). Brain atlas for glycoprotein hormone receptors at single-transcript level. ELife, 11, e79612. 10.7554/eLife.79612

Schumacher, A., & Zenclussen, A. C. (2019). Human Chorionic Gonadotropin-Mediated Immune Responses That Facilitate Embryo Implantation and Placentation. Frontiers in Immunology, 10. https://www.frontiersin.org/articles/10.3389/fimmu.2019.02896

Short, R. A., Bowen, R. L., O’Brien, P. C., & Graff-Radford, N. R. (2001). Elevated gonadotropin levels in patients with Alzheimer disease. Mayo Clinic Proceedings, 76(9), 906–909. 10.4065/76.9.906

Telegdy, G., Tanaka, M., & Schally, A. V. (2009). Effects of the LHRH antagonist Cetrorelix on the brain function in mice. Neuropeptides, 43(3), 229–234. 10.1016/j.npep.2009.03.001

Tsui. F.W., Martin, A., Wang, J., & Tsui, H.W. (2006). Investigations into the regulation and function of the SH2 domain-containing protein-tyrosine phosphatase, SHP-1. Immunol Res.;35(1-2):127–36. doi: 10.1385/IR:35:1:127. PMID: 17003515.

van Groenendael, R., Beunders, R., Kox, M., van Eijk, L. T., & Pickkers, P. (2019). The Human Chorionic Gonadotropin Derivate EA-230 Modulates the Immune Response and Exerts Renal Protective Properties: Therapeutic Potential in Humans. Seminars in Nephrology, 39(5), 496–504. 10.1016/j.semnephrol.2019.06.009

Verdile, G., Laws, S. M., Henley, D., Ames, D., Bush, A. I., Ellis, K. A., Faux, N. G., Gupta, V. B., Li, Q.-X., Masters, C. L., Pike, K. E., Rowe, C. C., Szoeke, C., Taddei, K., Villemagne, V. L., Martins, R. N., & AIBL Research Group. (2014). Associations between gonadotropins, testosterone and β amyloid in men at risk of Alzheimer’s disease. Molecular Psychiatry, 19(1), 69–75. 10.1038/mp.2012.147

Walton, J.C., Schilling, K., Nelson, R.J., & Oberdick, J. (2012). Sex-dependent behavioral functions of the Purkinje cell-specific Gαi/o binding protein, Pcp2(L7). Cerebellum. 11(4):982–1001. doi: 10.1007/s12311-012-0368-4.

Walker, C. K., & Herskowitz, J. H. (2021). Dendritic Spines: Mediators of Cognitive Resilience in Aging and Alzheimer’s Disease. The Neuroscientist : A Review Journal Bringing Neurobiology, Neurology and Psychiatry, 27(5), 487–505. 10.1177/1073858420945964

Woolley, C., & McEwen, B. (1992). Estradiol mediates fluctuation in hippocampal synapse density during the estrous cycle in the adult rat. The Journal of Neuroscience, 12(7), 2549–2554. 10.1523/JNEUROSCI.12-07-02549.1992

Xiong, J., He, C., Li, C., Tan, G., Li, J., Yu, Z., Hu, Z., & Chen, F. (2013). Changes of Dendritic Spine Density and Morphology in the Superficial Layers of the Medial Entorhinal Cortex Induced by Extremely Low-Frequency Magnetic Field Exposure. PLOS ONE, 8(12), e83561. 10.1371/journal.pone.0083561

Xiong, J., Kang, S. S., Wang, Z., Liu, X., Kuo, T.-C., Korkmaz, F., Padilla, A., Miyashita, S., Chan, P., Zhang, Z., Katsel, P., Burgess, J., Gumerova, A., Ievleva, K., Sant, D., Yu, S.-P., Muradova, V., Frolinger, T., Lizneva, D., … Ye, K. (2022). FSH blockade improves cognition in mice with Alzheimer’s disease. Nature, 603(7901), Article 7901. 10.1038/s41586-022-04463-0.

Yan J, Zhang Y, Yu H, Zong Y, Wang D, Zheng J, Jin L, Yu X, Liu C, Zhang Y, Jiang F, Zhang R, Fang X, Xu T, Li M, Di J, Lu Y, Ma X, Zhang J, Jia W, Hu C (2022). GPSM1 impairs metabolic homeostasis by controlling a pro-inflammatory pathway in macrophages. Nature 5;13(1), Article 7260. 10.1038/s41467-022-34998-9.

Yang, E.-J., Nasipak, B. T., & Kelley, D. B. (2007). Direct action of gonadotropin in brain integrates behavioral and reproductive functions. Proceedings of the National Academy of Sciences of the United States of America, 104(7), 2477–2482. 10.1073/pnas.0608391104

Zagrebelsky, M., Tacke, C., & Korte, M. (2020). BDNF signaling during the lifetime of dendritic spines. Cell and Tissue Research, 382(1), 185–199. 10.1007/s00441-020-03226-5

Zhang, W., Lei, Z. M., & Rao, C. V. (1999). Immortalized hippocampal cells contain functional luteinizing hormone/human chorionic gonadotropin receptors. Life Sciences, 65(20), 2083–2098. 10.1016/s0024-3205(99)00474-9

Ziegler, S. G., & Thornton, J. E. (2010). Low luteinizing hormone enhances spatial memory and has protective effects on memory loss in rats. Hormones and Behavior, 58(5), 705–713. 10.1016/j.yhbeh.2010.07.002

